# Deletion of Furin Facilitates Efficient and Versatile Coronavirus Pseudotyped Particle Generation

**DOI:** 10.64898/2026.11.19.767861

**Authors:** Pengcheng Wang, Yongtao He, Lianne Wu, Sam J. Wilson, Suzannah J. Rihn

## Abstract

Coronaviruses bring threats to both humans and animals. To initiate successful infections, coronaviruses must first navigate entry into a host cell. Coronavirus spike-bearing pseudotyped particle systems are indispensable tools for studying coronavirus entry and evaluating neutralizing antibodies and entry inhibitors. However, pseudotyped particle titers can be low when packaged particles bear coronavirus spikes that contain the furin cleavage site, and this has been a limitation for pseudotyped particle production and applications. Here, we demonstrate that CRISPR/Cas9-mediated knockout of furin protease in HEK293T cells significantly enhances the production of functional pseudotyped particles bearing spikes containing the furin cleavage site, including those from SARS-CoV-2, MERS-CoV, and MHV-JHM. Moreover, deletion of endogenous furin in producer cells yields an effect equivalent to deletion of the furin cleavage site, but with broader application across different spike proteins, and without virus manipulation. This work establishes a simple and effective platform to improve coronavirus pseudotyped particle systems by eliminating furin-mediated pre-activation, enabling more efficient and reliable production for downstream applications.

**IMPORTANCE:** The furin cleavage site (FCS) at the S1/S2 junction of coronavirus spike proteins is a critical determinant of viral infectivity, host tropism and transmissibility. Notably, the presence of an FCS in newly emerged SARS-CoV-2 was considered to be strongly associated with its efficient human-to-human spread and pandemicity. However, pseudotyped particles bearing FCS-containing spikes often exhibit low titers. This technical barrier hampers downstream applications such as neutralization assays, receptor identification, and entry mechanism studies. Here, we present a simplified and broadly applicable strategy that markedly improves pseudotyped particle yield by knocking out endogenous furin in producer cells. Our platform maintains spike integrity without requiring sequence modification and enables the robust production of functional pseudotyped particles for multiple FCS-containing coronaviruses, including SARS-CoV-2 and MERS-CoV. This work provides a practical and scalable tool for studying high-risk coronaviruses with pandemic potential.

## INTRODUCTION

Coronaviruses already pose numerous threats to humans and animals, and the potential for future zoonotic spillover of additional animal coronaviruses into humans further exacerbates this threat (1). In recent years, the SARS-CoV-2 pandemic has brought immense devastation to the world (2), and there is significant evidence indicating that SARS-CoV-2 emerged through a natural zoonotic spillover event (3–6). One defining feature of SARS-CoV-2 that may have promoted its pandemic potential is the presence of an unusual furin cleavage site (FCS) at the S1/S2 boundary of the SARS-CoV-2 spike protein (7). The FCS enables cleavage of spike by host furin proteases during viral egress, thereby pre-activating the spike for efficient entry into target cells (8). While the presence of an FCS is rare within the sarbecoviruses (expect SARS-CoV-2) (9), which is the subgenus SARS-CoV-2 belongs to, similar sites have been observed in other betacoronaviruses such as MERS-CoV, feline CoVs, and certain strains of murine hepatitis virus (MHV) (10–12).

MHV is a *Betacoronavirus*, belonging to the subgenus *Embecovirus*, that naturally infects mice and which utilizes murine CEACAM1 as its entry receptor (13). Among the various MHV strains, MHV-A59 and MHV-JHM are two well-characterized prototypes with distinct tissue tropisms and pathogeneses. MHV-A59 is hepatotropic and mildly neurotropic, while MHV-JHM is highly neurovirulent (14). Intriguingly, the spike protein of MHV-JHM can mediate cell receptor-independent infection and spread in the central nervous systems of *Ceacam1*^-/-^ mice (15,16). MHV-JHM also has been reported to infect a human hepatocellular carcinoma cell line (17). These findings raise interest in whether it may utilize alternative receptors and prompt concerns about its potential to spill over into humans.

To enable safe and quantitative studies of coronavirus entry under biosafety level 2 (BSL-2) conditions, researchers have developed diverse pseudotyped particle-based systems, which generate non-replicating virus-like particles that are engineered to express the coronavirus spike protein (18,19). These systems can be used to ascertain or validate virus receptors, identify entry inhibitors, and assess the neutralizing activity of antibodies (19,20). Typically, pseudotyped particles systems are constructed using HIV-, VSV- or MLV-derived platforms and often employ reporter genes such as GFP, NanoLuc, or firefly luciferase to enable quantification (21). As there has been a strong need for pseudotyped particle systems to evaluate and detect SARS-CoV-2 therapeutics and neutralization, numerous strategies have been developed to improve yields of SARS-CoV-2 spike-pseudotyped particles. These include truncation of the spike cytoplasmic tail (22,23), deletion of the spike furin cleavage site (24), optimization of plasmid ratios (25) and diverse concentration protocols (26,27).

In this investigation, we initially set out to investigate the host tropism of the MHV-JHM strain, following reports, as mentioned above, that its entry may not be species- or CEACAM1-dependent. To do this, we generated pseudotyped lentiviral vector-based particles bearing the MHV-JHM spike protein; however, these particles yielded a low viral titer. We observed pronounced cell–cell fusion during the production process, leading us to hypothesize that perhaps the S1 subunit had already dissociated during particle production, thereby exposing the fusion peptide. We therefore developed a furin-deficient producer cell system to improve the production of pseudotyped particles bearing furin cleavage site. This system overcomes common limitations in pseudotyping cleavable spikes and facilitates safe downstream applications in viral entry studies, neutralization assays, and antiviral screening.

## RESULTS

### Furin-deficiency increases MHV-JHM functional pseudotyped particle production

MHV-JHM, unlike other MHVs, has been reported to mediate CEACAM1-independent cell entry and exhibit the potential to infect human cell lines (15,16). To further elucidate the determinants of MHV-JHM entry and investigate its species specificity, we generated pseudotyped lentiviral particles bearing the spike proteins of MHV-JHM and MHV-A59. Surprisingly, the MHV-JHM-pseudotyped particles exhibited extremely low titers in HEK293T cells (Fig. 1A), in contrast to particles pseudotyped with MHV-A59. In addition, we observed extensive syncytia formation during particle production in cells expressing the MHV-JHM spike (Fig. 1B), suggesting that the MHV-JHM spike protein may undergo premature activation due to furin-mediated cleavage, resulting in exposure of the fusion peptide. To test whether proteolytic processing was responsible for the low titer we observed, we targeted FURIN, transmembrane protease serine 2 (TMPRSS2), and cathepsin L (CTSL) individually using CRISPR/Cas9. Furin deficiency was confirmed by western blotting, and CTSL targeting resulted in partial depletion of the protein (Supplementary Fig. 1A). For TMPRSS2, because its endogenous protein expression was too low for reliable detection, the CRISPR editing efficiency was assessed at the genomic level using Sanger sequencing and Inference of CRISPR Edits (ICE) analysis (28) (Supplementary Fig. 1B). Among these three, furin deficiency remarkably increased MHV-JHM-pseudotyped particle titers by approximately 30-fold (or 1.5 logs) (Fig. 1C), whereas targeting TMPRSS2 or CTSL had only very modest effects on pseudotyped particle titers (<2-fold). Consistent with these findings, treating producer cells with a furin inhibitor, Decanoyl-RVKR-CMK, during pseudotyped particle production also markedly enhanced MHV-JHM-pseudotyped particle titers (Fig. 1E). In contrast to these results, neither genetic deficiency nor pharmacological inhibition of furin improved the titers of MHV-A59-pseudotyped particles (Fig. 1D and 1F). We confirmed furin knockout by western blot and assessed knockout efficiency by Sanger sequencing and ICE analysis, and established clonal furin knockout cell lines (Fig. 1G and 1H). Because HEK293T cells are aneuploid and pseudotriploid, Sanger sequencing alone did not unambiguously resolve the inserted nucleotide sequences at the edited furin locus. We therefore performed Oxford Nanopore long-read sequencing and analyzed the resulting high-throughput sequencing data using CRISPResso (29) to characterize the inserted nucleotide sequences of clone 7 (Supplementary Fig. 1C), which we used as our furin KO cells for the rest of this study. In subsequent analyses, furin knockout did not affect the production of VSV-G or MHV-A59-pseudotyped particles but markedly increased the titer of MHV-JHM-pseudotyped particles (Fig. 1I). We further targeted TMPRSS2 or CTSL in the monoclonal furin knockout cell line by CRISPR-Cas9 and found that this did not further increase the titer of the MHV-JHM-pseudotyped particles (Fig. 1J), and this may be because furin acts as the upstream protease during spike maturation. Taken together, these results demonstrate the establishment of an efficient packaging platform for MHV-JHM-pseudotyped particles.

**Figure 1.**
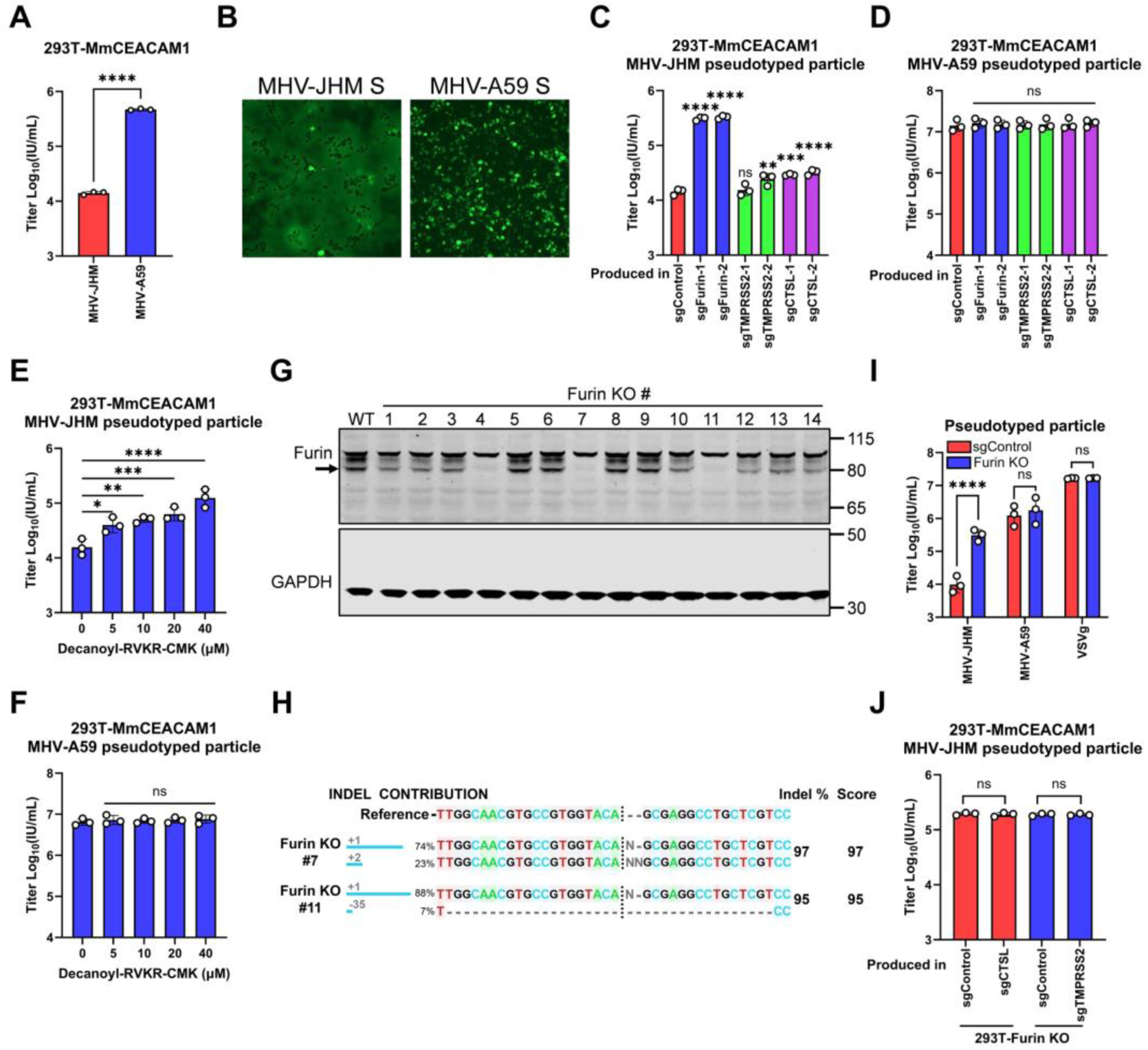
Furin deficiency increases MHV-JHM functional pseudotyped particle titers. **A.** Comparison of titers of HIV-based pseudotyped particles bearing MHV-JHM or MHV-A59 spike proteins produced in HEK293T cells on 293T-MmCEACAM1 cells. **B.** Representative fluorescence images of HEK293T cells transfected with MHV-JHM or MHV-A59 spike expression plasmids during pseudotyped particle production. **C-D**. MHV-JHM (C) and MHV-A59 (D) pseudotyped particles were produced in control or protease-targeted HEK293T cells and titrated on 293T-MmCEACAM1 cells. **E-F.** MHV-JHM (E) and MHV-A59 (F) pseudotyped particles were produced in the presence of Decanoyl-RVKR-CMK or DMSO, and infectious titers were measured on 293T-MmCEACAM1 cells. **G.** Following lentiviral transduction and antibiotic selection, 293T-sgFurin cells were subjected to limiting dilution to isolate monoclonal cell lines, which were expanded and analyzed by immunoblotting for furin expression, with GAPDH as a loading control. **H.** Single colonies were expanded for genomic DNA extraction. The sgRNA target regions were amplified, and knockout efficiency was determined by ICE analysis. **I.** Titers of pseudotyped particles bearing MHV-JHM spike, MHV-A59 spike, or VSV-G envelope produced in control or furin-knockout HEK293T cells. **J.** Infectious titers of MHV-JHM pseudotyped particles produced in furin/CTSL or furin/TMPRSS2-edited HEK293T cells and measured on 293T-MmCEACAM1 cells. Data shown are from three independent experiments, and each independent experiment was performed in triplicate. (A) Unpaired t test; ( C, D, E, and F) one-way ANOVA with Dunnett’s test; (I and J) two-way ANOVA with Sidak’s test; mean ± s.d.; \**P* < 0.05; \*\**P* < 0.01; \*\*\**P* < 0.001; \*\*\*\**P* < 0.0001; ns, not significant.

### Knockout of furin decreases cell fusion induced by the MHV-JHM spike while maintaining its integrity

To elucidate the role of furin in spike-induced cell–cell fusion during MHV-JHM-pseudotyped particle production, we established a split GFP/NanoLuc–based cell–cell fusion assay (Fig. 2A). By direct fluorescence imaging, quantification of GFP fluorescence, and NanoLuc luminescence measurements, we found that MHV-JHM, but not A59, induces *Mus musculus* CEACAM1 (MmCEACAM1)-independent cell–cell fusion (Fig. 2B-D). Moreover, upon furin knockout, transfection of the MHV-JHM spike induced markedly reduced cell–cell fusion, indicating that furin deletion reduces fusion peptide exposure in MHV-JHM spike. (Fig. 2E-G). To clarify how furin knockout increases the infectious titer of MHV-JHM pseudotyped particles, we first confirmed by qPCR that deletion of furin in producer cells does not alter the viral genome copy number of MHV-JHM pseudotyped particles. Consistently, the level of pelletable p24 protein in the pseudotyped particle-containing supernatant was also comparable between groups (Fig. 2H). Furthermore, we examined spike protein expression following transfection in the presence or absence of furin. Using an MHV S1-specific antibody, we detected a greater abundance of full-length MHV-JHM spike in furin-deficient cells, and detection through a C-terminal HA tag on spike revealed a higher mobility spike product in control cells. In contrast, the MHV-A59 spike remained full-length regardless of furin expression (Fig. 2I). Following sucrose cushion centrifugation of the pseudotyped particle-containing supernatants, western blot analysis revealed markedly increased incorporation of full-length MHV-JHM spike into particles produced in furin-knockout cells. By contrast, full-length spike was barely detectable in particles generated in control cells. (Fig. 2J). Together, these results support a model in which furin knockout limits the premature proteolytic processing of MHV-JHM spike, preserves the incorporation of full-length spike into pseudotyped particles, and may help maintain the spike in a prefusion state, thereby reducing spike-mediated cell–cell fusion during particle production.

**Figure 2.**
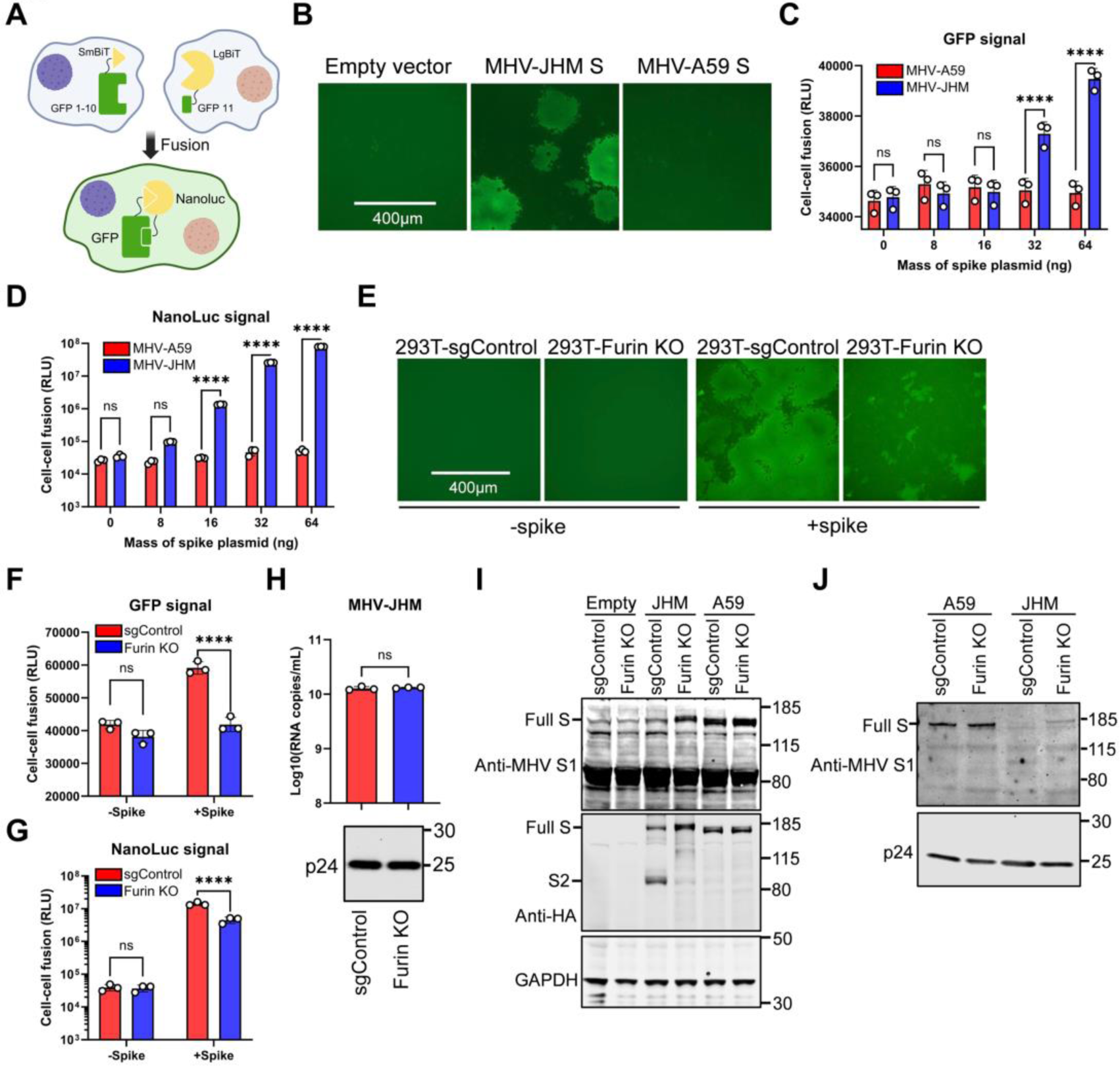
Knockout of furin decreases cell fusion induced by MHV-JHM spike and maintains its integrity. **A.** Schematic of cell-cell fusion assay. **B-D.** 293T-GFP1-10-SmBiT and 293T-GFP11-LgBiT were mixed and after transfection with MHV-JHM or MHV-A59 spike protein, spike protein-induced syncytia were visualized under a fluorescence microscope (B) or quantified by GFP (C) and NanoLuc (D) signal. **E-G.** 293T-sgControl-GFP1-10-SmBiT cells were mixed with 293T-sgControl-GFP11-LgBiT cells, or 293T-furin-knockout-GFP1-10-SmBiT cells were mixed with 293T-furin-knockout-GFP11-LgBiT cells, followed by transfection with MHV-JHM spike or empty vector. spike protein-induced syncytia were visualized under a fluorescence microscope (E) or quantified by GFP (F) and NanoLuc (G) signal. **H.** MHV-JHM pseudotyped particles were produced in control and furin-knockout cells. Supernatants were collected at 48 h post-transfection, followed by RNA extraction and reverse transcription. Pseudotyped particle copy numbers were then quantified by absolute RT-qPCR using GFP-specific primers. In parallel, pseudoviral particles were concentrated through a sucrose cushion by centrifugation and analyzed by western blotting for HIV-1 p24 to assess particle production. **I.** S protein expression levels were assessed in cells lysed by western blot using antibodies against MHV-S1, the spike C-terminal HA tag and GAPDH as internal control. The full-length spike (Full S) and S2 subunit (S2) are indicated. **J.** Pseudotyped particles produced in control or furin-knockout HEK293T cells were pelleted through a layer of 20% of sucrose, followed by western blot using antibodies against MHV-S1 and HIV p24 as control. Data shown are from three independent experiments, and each independent experiment was performed in triplicate. (C, D, F, and G) two-way ANOVA with Sidak’s test; (H) Unpaired t test; mean ± s.d.; \**P* < 0.05; \*\**P* < 0.01; \*\*\**P* < 0.001; \*\*\*\**P* < 0.0001; ns, not significant.

### Parental MHV-JHM-pseudotyped particles fail to enter human liver carcinoma cells

Using our platform, we next produced high-titer MHV-JHM pseudotyped particles and assessed their ability to enter human Huh7 cells. While human seasonal coronavirus HCoV-229E-pseudotyped particles infected Huh7 cells with high efficiency, high-titer MHV-JHM-pseudotyped particles failed to enter these cells (Fig. 3A). We examined previously reported MHV-JHM strains and found that only a derivative strain known as MHV-JHM-DL, which was obtained through serial passage of MHV-JHM in a suckling mouse brain, was capable of entering Huh7 cells (30). Based on the limited publicly available sequencing information for MHV-JHM-DL, we replaced a ∼1,000 bp region of the MHV-JHM genome with the corresponding sequence from MHV-JHM-DL and generated pseudotyped particles bearing this chimeric spike, termed MHV-JHM-DL-R1K, using our same furin knockout platform (Fig. 3B). The results showed that the chimeric MHV-JHM-DL-R1K-pseudotyped particles were also unable to enter Huh7 cells. Moreover, the entry efficiency of the MHV-JHM-DL-R1K-pseudotyped particles into MmCEACAM1-expressing cells was markedly reduced, and furin knockout also failed to enhance the titer of MHV-JHM-DL-R1K pseudotyped particle. These findings suggest that replacement of only a ∼1,000 bp region generates a functionally defective chimeric spike protein and parental MHV-JHM pseudotyped particle is not permissive for entry into human liver carcinoma cells.

**Figure 3.**
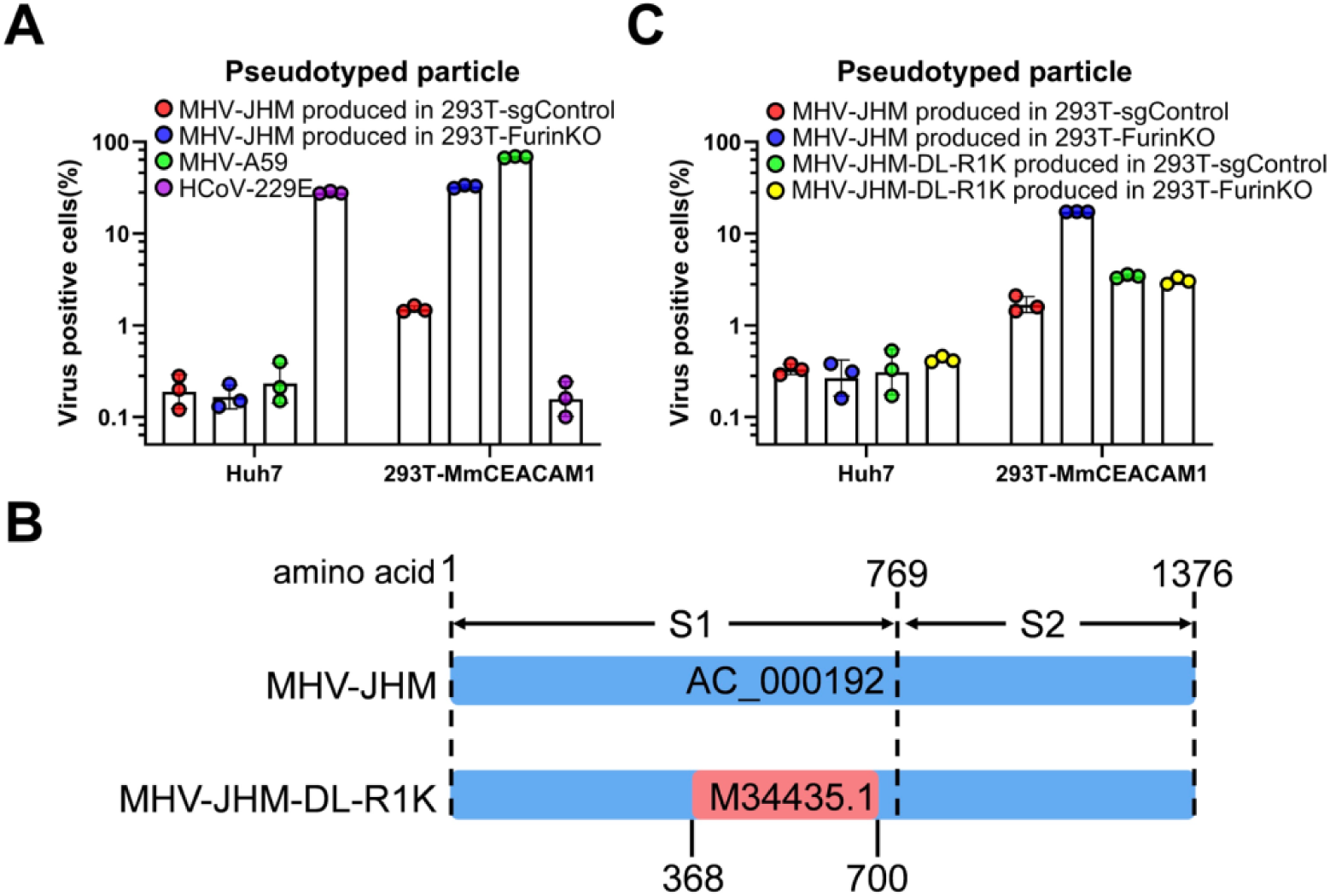
Parental MHV-JHM pseudotyped particles fail to enter human liver carcinoma cells. **A.** Parental MHV-JHM pseudotyped particles were produced in control or furin-knockout HEK293T cells, whereas MHV-A59 and HCoV-229E pseudotyped particles were produced in wild-type HEK293T cells. These pseudotyped particles (100 µL) were then used to infect Huh7 or 293T-MmCEACAM1 cells in 96-well plates, and GFP-positive cells were quantified by flow cytometry at 48 h postinfection. **B.** Schematic of the MHV-JHM and MHV-JHM-DL-R1K spike sequences, in which an approximately 1,000-bp published MHV-JHM-DL sequence was substituted into the parental MHV-JHM spike at the C-terminal domain of S1. **C.** Parental MHV-JHM and chimeric MHV-JHM-DL-R1K pseudotyped particles were produced in control or furin-knockout HEK293T cells. These pseudotyped particles (100 µL) were used to infect Huh7 or 293T-MmCEACAM1 cells in 96-well plates, and GFP-positive cells were quantified by flow cytometry at 48 h postinfection. Data shown are from three independent experiments, and each independent experiment was performed in triplicate.

### Furin knockout enhances titers of SARS-CoV-2- and MERS-CoV-pseudotyped particles

To evaluate the applicability of our platform to highly pathogenic human coronaviruses, we next produced SARS-CoV-2- and MERS-CoV-pseudotyped particles using furin-knockout producer cells. Furin deficiency increased SARS-CoV-2 and MERS-CoV pseudotype titers by approximately 15.6- and 4.7-fold respectively (Fig. 4A). The relatively modest increase in MERS-CoV pseudotyped particles titer may be attributable to the lower susceptibility of its furin cleavage site to furin-mediated processing (31). Given the strong cell–cell fusion induced by MHV-JHM spike, we further evaluated whether the SARS-CoV-2 or MERS-CoV spikes could also mediate cell–cell fusion. We therefore mixed 293T cells only stably expressing split GFP/NanoLuc fragments and transfected them with SARS-CoV-2 or MERS-CoV spikes. Using the highly sensitive NanoLuc readout, we did not detect any cell–cell fusion, indicating that the fusion peptide was not exposed (Fig. 4B). To determine whether SARS-CoV-2 spike proteins were pre-cleaved during production, we analyzed spike expression by western blot. In furin-knockout cells, the SARS-CoV-2 spike was detected predominantly in its full-length form, whereas cleaved S1 fragments were observed in control cells(Fig. 4C). After purifying pseudotyped particles through sucrose cushion centrifugation, western blot analysis showed significantly higher levels of full-length SARS-CoV-2 spike protein in particles from furin-knockout cells compared to control cells, where the full-length spike was barely detected (Fig. 4D). Overall, similar to MHV-JHM pseudotyped particle production, SARS-CoV-2 spike was pre-activated in control 293T cells. However, the difference is that SARS-CoV-2 spike did not expose its fusion peptide. Furin knockout of producer cells enhances intact S protein incorporation into SARS-CoV-2 pseudotyped particles.

**Figure 4.**
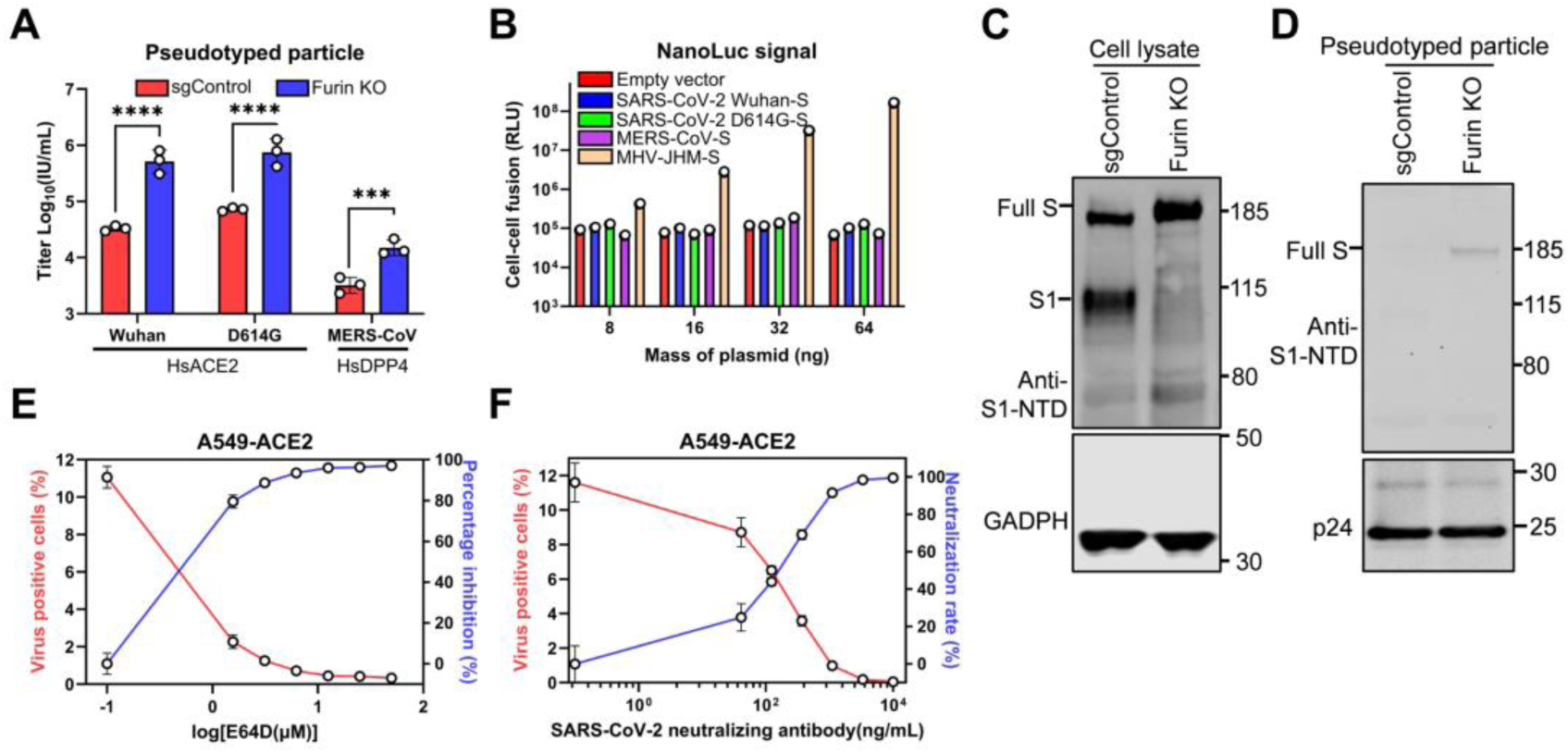
Furin knockout enables efficient packaging of SARS-CoV-2 and MERS-CoV pseudotyped particles. **A.** SARS-CoV-2 (original Wuhan-1 strain and D614G variant) and MERS-CoV pseudotyped particles were produced in control or furin-knockout HEK293T cells and titrated on cells exogenously expressing the corresponding receptors. **B.** 293T-GFP1-10-SmBiT and 293T-GFP11-LgBiT were mixed and transfected with SARS-CoV-2, MERS-CoV or MHV-JHM spike proteins, and syncytia were then quantified by NanoLuc signal. **C.** Western blot analysis of the expression and processing of SARS-CoV-2 spike in the pseudotyped particles producer cells at 48 h after transfection with antibodies against SARS-CoV-2 S1. GAPDH was used as a loading control for the cell lysate. Full-length spike (Full S) and SARS-CoV-2 S1 subunit (S1) are indicated. **D.** SARS-CoV-2 pseudotyped particles produced in control or furin-knockout HEK293T cells were pelleted through a layer of 20% of sucrose, followed by western blotting using antibodies against SARS-CoV-2 S1. Pelletable HIV p24 was used as a loading control for pseudotyped particles. **E.** A549-ACE2 cells were pretreated with E64D and infected with 100 µL of SARS-CoV-2 pseudotyped particles produced in furin-knockout HEK293T cells. GFP-positive cells were quantified by flow cytometry at 48 h postinfection. The percentage inhibition was calculated as follows: percentage of inhibition (%) = 100 × (1 – GFP-positive cells rate of E64D treatment / GFP-positive cells rate of DMSO treatment). **F.** Using 100 µL of SARS-CoV-2 pseudotyped particles produced in furin-knockout 293T cells, the pseudotyped particles was incubated with the indicated concentration of neutralizing antibody at 37 °C for 1 h. The original medium was then removed from A549-ACE2 cells, and the pseudotyped particles mixture was added. GFP-positive cells were quantified by flow cytometry at 48 h postinfection. The neutralization rate was calculated as follows: percentage of neutralization (%) = 100 × (1 – GFP-positive cells rate of antibody treatment / GFP-positive cells rate of DMEM treatment). Data shown are from three independent experiments, and each independent experiment was performed in triplicate. (A) Two-way ANOVA with Sidak’s test; mean ± s.d.; \**P* < 0.05; \*\**P* < 0.01; \*\*\**P* < 0.001; \*\*\*\**P* < 0.0001; ns, not significant.

We next utilized high-titer SARS-CoV-2 pseudotyped particles in an entry pathway study. A549 cells exogenously expressing ACE2 (A549-ACE2) cells were pretreated with DMSO or different concentrations of E64D, an inhibitor of endosomal entry, and subsequently infected with pseudotyped particles produced in furin knockout 293T cells. We found that high-titer pseudotyped particles yielded stable and reproducible inhibition curves (Fig. E). Neutralization can be quantified by incubating antibody with high-titer pseudotyped particles at 37 °C for 1 h, and then adding the mixture to A549-ACE2 cells (Fig. 4F). Our system is equally effective for SARS-CoV-2 and MERS-CoV, and high-titer pseudotyped particles can be used to study viral entry and neutralizing antibodies.

### Knockout of endogenous furin provides broader utility than deletion of FCS for spikes with FCS

Previous studies have shown that deletion of the FCS in SARS-CoV-2 spikes can be used to increase pseudotyped particle titers or to stabilize spike trimers for structural studies (24,32,33). We therefore wanted to directly compare the effects and applicability of endogenous furin knockout versus FCS removal as two distinct strategies for improving pseudotyped particle production. We mutated the furin cleavage sites of MHV-JHM, SARS-CoV-2, and MERS-CoV spikes to GSAS, and conversely replaced an active RRAR motif at the S1/S2 junction of MHV-A59, and then packaged the pseudotyped particles using either control or furin-knockout HEK293T cells (Fig. 5A). The results showed that for SARS-CoV-2, endogenous furin knockout and FCS removal had comparable effects, each increasing pseudotyped particle titers by approximately 16-fold relative to wild-type SARS-CoV-2 pseudotyped particle produced in control cells (Fig. 5C). In contrast, for MHV-JHM and MERS-CoV, furin knockout was more effective at raising particle titers than FCS removal. Notably, for MERS-CoV, deletion of the FCS in control cells even resulted in a reduction in pseudotyped particle titer (Fig. 5B and 5D). For MHV-A59, which naturally lacks an active FCS, introduction of an RRAR motif into the spike protein reduced pseudotyped particle titers; however, this reduction was reversed when the virus was produced in furin-knockout cells (Fig. 5E). Collectively, these findings indicate that endogenous furin knockout offers broader applicability than FCS removal for improving pseudotyped particle production. To determine whether the inappropriately cleaved WT spike exerted a dominant-negative effect on the uncleaved GSAS-spike, we produced pseudotyped particles in control 293T cells by co-transfecting the WT and GSAS spike at different ratios and subsequently measured their infectious titers. The pseudovirus titer decreased approximately linearly as the proportion of WT spike increased, indicating no detectable dominant-negative effect of the WT spike on the GSAS-mutant spike (Supplementary Fig. 2A and B).

**Figure 5.**
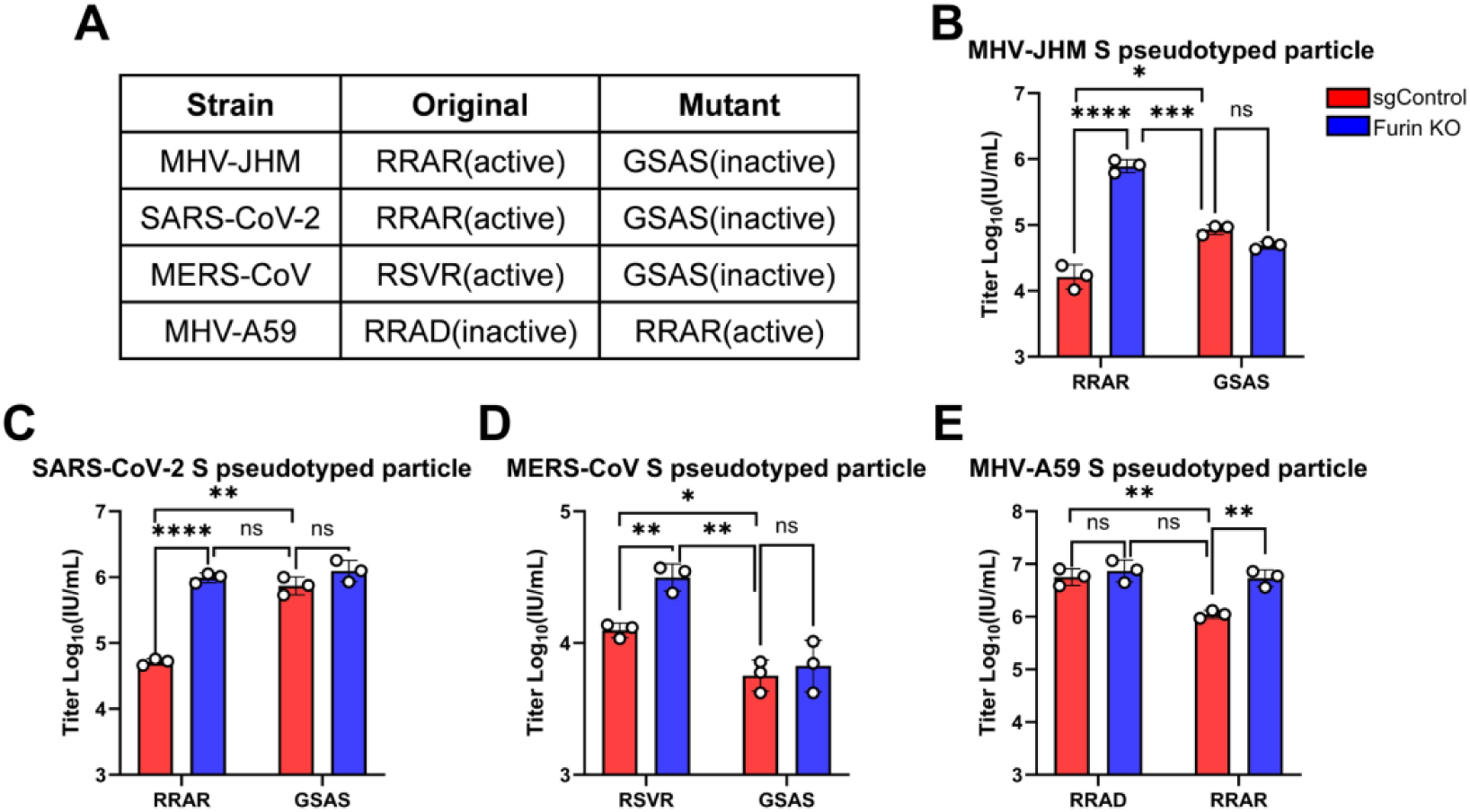
Knockout of endogenous furin provides broader utility than deletion of FCS for spikes with FCS. **A.** Summary of FCS mutations introduced into spike proteins from MHV-JHM, SARS-CoV-2, MERS-CoV, and MHV-A59. **B-E.** Pseudotyped particles bearing wild-type or mutant spike proteins of MHV-JHM (B), SARS-CoV-2 (C), MERS-CoV (D), and MHV-A59 (E) were produced in control or furin-knockout HEK293T cells and titrated on HEK293T cells stably expressing the corresponding receptors. Data shown are from three independent experiments, and each independent experiment was performed in triplicate. (B, C, D, and E) Two-way ANOVA with Sidak’s test and Unpaired t test; mean ± s.d.; \**P* < 0.05; \*\**P* < 0.01; \*\*\**P* < 0.001; \*\*\*\**P* < 0.0001; ns, not significant.

In summary, our findings suggest that betacoronaviruses, like MHV-A59, without an active FCS can achieve high pseudotyped particle titers in conventional producer cells. In contrast, for betacoronaviruses like MHV-JHM and SARS-CoV-2, the presence of a potent FCS leads to furin-mediated pre-cleavage of the spike, resulting in partial loss of the S1 subunit and reduced functional pseudotyped particle production. In the case of MHV-JHM, premature cleavage further exposes the fusion peptide, triggering extensive cell–cell fusion. For coronaviruses bearing an FCS, deletion of endogenous furin in producer cells markedly enhances pseudotyped particle titers (Fig. 6).

**Figure 6.**
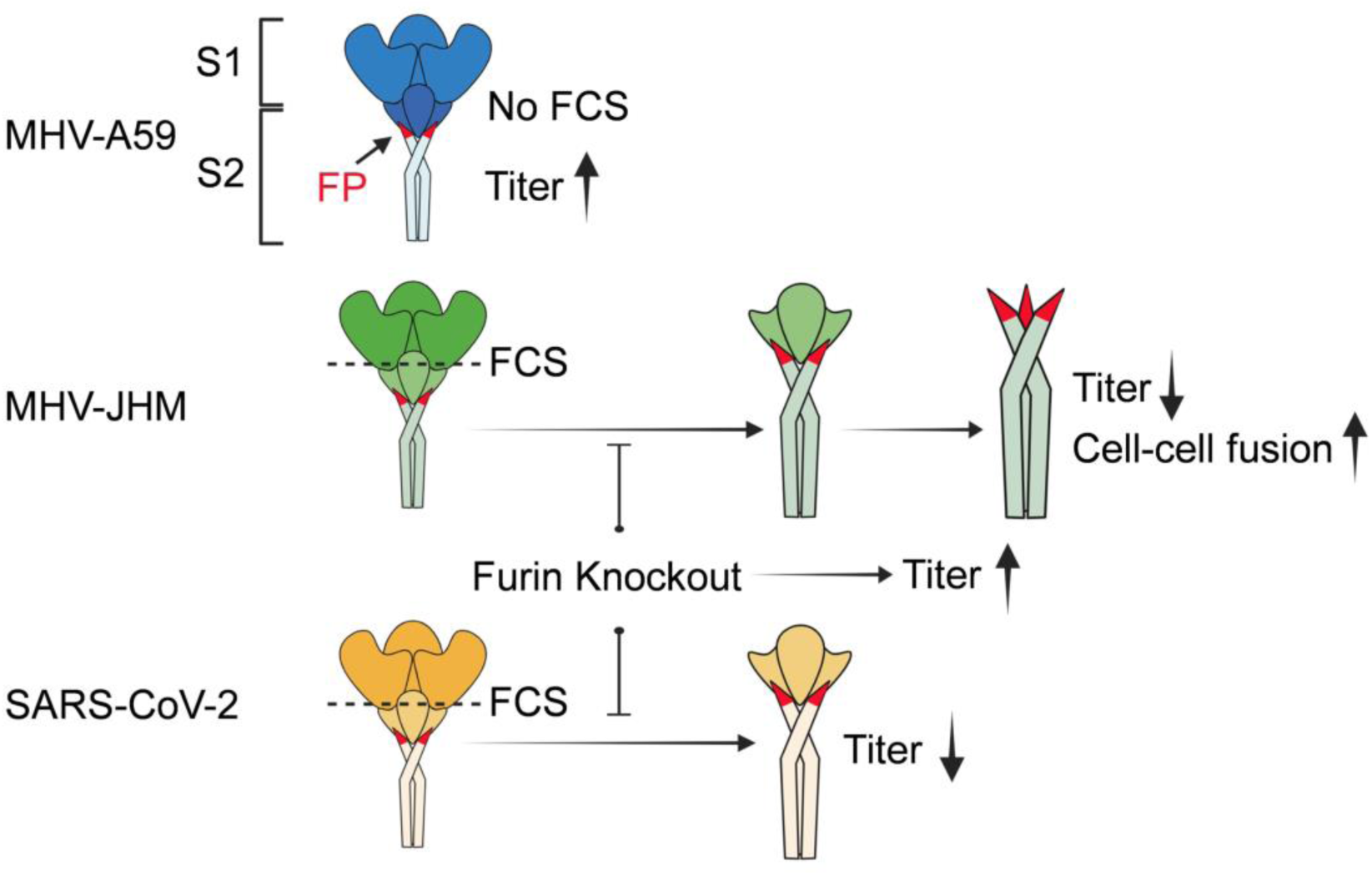
Graphic summary. Schematic model illustrating the effects of endogenous furin on cell – cell fusion and pseudotyped particle production for coronaviruses. FP, fusion peptide; FCS, furin cleavage site. Illustration was partially created with BioRender.com.

## DISCUSSION

In this study, we successfully established a high efficiency coronavirus pseudotyped particle platform by knocking out the endogenous furin protease. This approach enhanced intact S protein incorporation into pseudotyped particles and significantly increased the pseudotyped particle titers of coronaviruses bearing an FCS. Compared to direct FCS removal, furin knockout offers broader applicability and operational simplicity.

In this study, we found that during the production of SARS-CoV-2 and MHV-JHM spike-pseudotyped particles in wild-type HEK293T cells, spike proteins underwent substantial proteolytic cleavage. Distinct S1/S2 cleavage products were readily detected, whereas the abundance of full-length spike was markedly reduced. Moreover, spike incorporation into the pseudoviral particles was barely detectable. In contrast, deletion of furin in the producer cells largely abolished spike cleavage into S1 and S2. Consequently, intact full-length spike was readily detected in the pseudoviral particles generated in the absence of furin, and this was accompanied by a marked increase in infectious titer. A similar phenotype has been reported following truncation of the spike cytoplasmic tail, which stabilizes the S1–S2 association and promotes more efficient incorporation of spike into SARS-CoV-2 pseudotyped particles (34). To overcome the instability and proteolytic cleavage of MHV-JHM spike during pseudovirus production, previous studies employed an alternative approach without modifying endogenous furin, generating virus-like particles (VLPs) containing the MHV-JHM spike, envelope (E), and membrane (M) proteins to facilitate stable spike incorporation (35). We speculate that M and E induce the retention of spike in ERGIC or cis-Golgi and reduce the amount of unassembled spike that proceeds toward the trans-Golgi network and plasma membrane (36). This may limit spike exposure to downstream proteases, including furin (37), and prevent excessive cleavage or degradation. Our approach, which directly removes furin from the producer cells, may achieve a similar outcome through a distinct mechanism.

During the production of authentic SARS-CoV-2, an appropriate pre-activation of spike by furin may not only facilitate the adoption of an open, receptor-binding-competent form, but also prime spike for subsequent membrane fusion (38). Consistent with this notion, furin deficiency has been reported to impair the propagation of authentic SARS-CoV-2 (39,40). However, our observations indicate that knockout of furin in producer cells significantly enhances the titer of SARS-CoV-2 pseudotyped particles. We speculate that when spike is expressed in the absence of other coronavirus structural proteins (as is the case in pseudotyped systems where spike is incorporated into the membrane surrounding a heterologous viral core), that furin-mediated S1/S2 cleavage may destabilize the non-covalent association between S1 and S2, promote premature S1 shedding, and reduce the incorporation of the complete spike into pseudotyped particles.

Using pseudotyped particles generated from our platform, we found that both parental MHV-JHM and a chimeric variant carrying a 1,000 bp spike fragment from the reported derivative strain failed to enter Huh7 cells. The inability of the parental MHV-JHM strain to infect Huh7 cells is expected, as no previous study has reported such tropism. The MHV-JHM-DL strain, which was reported to infect Huh7 cells, was derived through serial passaging of MHV-JHM in suckling mouse brains, possibly altering its host tropism during the process (30). However, only a 1,000 bp segment of the MHV-JHM-DL spike sequence is publicly available. We substituted this region into the parental spike to generate a chimeric spike MHV-JHM-DL-R1K, but the chimeric spike pseudotyped particles still failed to enter Huh7 cells and even showed reduced entry into cells expressing mouse CEACAM1. This suggests that the artificial chimeric spike may not faithfully reproduce the function of the authentic MHV-JHM-DL spike protein. Therefore, our results cannot conclusively reflect the entry capacity of the actual MHV-JHM-DL strain.

Removal of the FCS is a common approach used to prevent conformational change to the post-fusion state during spike protein expression, and in particular, is used to facilitate crystallography studies (41,42). In addition, FCS deletion has also been used to enhance the production of SARS-CoV-2 pseudotyped particles (24). Our study demonstrates that, for pseudotyped particle production, knockout of endogenous furin offers a broader applicability than FCS removal, without adjustment to the spike sequence. We speculate that our platform may also be advantageous for expressing stable spike proteins, potentially benefiting structural and biophysics studies.

At present, numerous SARS-CoV-2 pseudotyped particle systems, particularly those based on the VSV platform, have been successfully employed in studies of viral neutralizing antibodies, entry pathways, and receptor identification (43–45). In this study, we developed a high-efficiency packaging platform based on the HIV pseudotyped particle system. This system uses a simple three-plasmid co-transfection strategy, offering a more streamlined and cost-effective workflow.

In summary, we have established a convenient and efficient pseudotyped particle production platform for coronaviruses bearing FCS by knocking out endogenous furin protease in producer cells. This approach enables the generation of high-titer pseudotyped particles without the need for viral sequence modification or additional concentration steps. Our platform offers advantages in simplicity, broad applicability, and cost, making it a valuable tool for studies of coronavirus entry, neutralizing antibodies, and antiviral screening.

## MATERIALS AND METHODS

### Cells

HEK293T, A549 and Huh7 cells were cultured in Dulbecco’s Modified Eagle Medium (DMEM) supplemented with 9% fetal bovine serum (FBS) and 50 mg/ml gentamicin and grown at 37 °C in 5% CO_2_. All cell lines were tested routinely and were free of mycoplasma contamination.

### Plasmids construction

The full-length sequence of MHV-JHM (YP_209233.1), MHV-A59 (NP_045300.1), SARS-CoV-2 (QHD43416.1), and MERS-CoV (YP_009047204.1) spikes were codon-optimized (except MERS-CoV) and synthesized by Integrated DNA Technologies company. The fragments were subsequently cloned into the pcDNA3.1(+) expression vector using Gibson assembly (NEB #E2611). For the MHV-JHM-DL-R1K spike construct, the published partial spike sequence of MHV-JHM-DL (M34435.1) was synthesized, and the remaining part of the spike gene was derived from MHV-JHM. The three fragments were fused by 2-round splicing by overlap extension (SOE)-PCR (NEB # M0492) to generate the full-length chimeric spike, which was subsequently cloned into the pcDNA3.1(+) vector using Gibson assembly. For detection of spike expression by western blot on Figure 2I, full-length MHV-JHM and MHV-A59 spike proteins were fused at the C-terminus to a GGGS linker followed by an HA tag. Removal or insertion of the FCS motif in spike proteins was performed using SOE-PCR, followed by cloning into the pcDNA3.1 expression vector via Gibson assembly. The lentiviral plasmids expressing human ACE2 (NP_001358344.1), human DPP4 (NM_001935.4) and mouse *Ceacam1* (NM_001039186.1) genes were synthesised, amplified, and cloned into the pLV-EF1α-IRES-hygro vector (Addgene #85134) using Gibson assembly.

The GFP and NanoLuc domains used for the split GFP/NanoLuc fusion system were PCR amplified from CSGW (46) and pCCI-4K-SARS-CoV-2-NanoLuc (MT926412.1) plasmids. The PCR products were then assembled into the pLV-EF1α-IRES-blast vector using Gibson assembly. A flexible 3 × GGGS linker was introduced between fusion elements. Finally, two lentiviral plasmids expressing GFP1-10-SmBiT and GFP11-LgBiT, respectively, were constructed. All plasmid sequences were verified using Oxford Nanopore Technology by Plasmidsaurus company.

### Pseudotyped particle production and infectivity assays

HIV-based pseudotyped particles were produced in HEK293T cells. Pseudotyped particle were produced by co-transfection of 5 μg CSGW reporter lentiviral vector that encodes GFP, 5 μg NL4.3 GagPol, and pcDNA3.1(+) vector expressing the spike of MHV spike (2 μg), SARS-CoV-2 (8 μg) or MERS-CoV (8 μg) using polyethylenimine (PEI). After 48 h, the pseudotyped particle particles were harvested and passed through 0.45 μm filters, and then aliquoted for storage at −80 °C. Pseudotyped particle titration was performed on HEK293T cells stably overexpressing mouse CEACAM1, human ACE2, or human DPP4 for MHV, SARS-CoV-2, and MERS-CoV pseudotyped particles, respectively. Pseudotyped particles were serially diluted 3-fold in cell culture medium and 100 μL of each dilution was overlaid onto the target cells in 96-well plates. After 48 hours of infection, cells were resuspended and fixed with 4% paraformaldehyde (PFA), followed by flow cytometry analysis (Cytek Guava) to determine the percentage of GFP-positive cells.

The lentiviruses used to exogenously express CEACAM1, ACE2 and DPP4, or to knockdown protein expression, were produced by co-transfecting HEK293T cells with 5 μg of the pLV-EF1α-IRES-hygro expressing the individuals receptors (or LentiCRISPR V2 vector), 5 μg of a lentiviral transfer and packaging plasmid (NL4.3 GagPol), and 1 μg of VSVg (glycoprotein from vesicular stomatitis virus) using polyethylenimine (PEI) in serum-free DMEM. Viral supernatant was collected at 48 hours post-transfection and filtered through 0.45 μm filters, and the filtered supernatant was then used to transduce HEK293T cells by spinoculation (1 hour, 1600 rpm, room temperature) with 6 μg/ml polybrene.

### Cell line generation and single colony

Modified HEK293T cell lines were generated by lentiviral transduction (pLV-EF1α-IRES-hygro) using vectors encoding mouse CEACAM1, human ACE2, human DPP4, or split GFP/NanoLuc subunits and selected with antibiotic for 7 days. The furin-deficiency HEK293T cell lines were generated by transducing packaged lentiCRISPR v2 Puro (Addgene #98290) expressing representative sgRNA. TMPRSS2- and CTSL-targeted HEK293T cells were generated by transducing packaged lentiCRISPR v2 Hygro expressing representative sgRNA. To establish a double-targeted cell line, furin-knockout HEK293T clonal cell lines were obtained by limiting dilution and verified by western blot and Inference of CRISPR Edits (ICE) analysis, and further transducing lentiviruses expressing individual sgRNA target TMPRSS2 or CTSL. After transducing, the cell lines were selected with antibiotic for 7 days. sgRNA sequences used for CRISPR editing are listed in Supplementary Table 1.

### Cell-cell fusion assay

Equal numbers of HEK293T or furin knockout cells stably expressing either GFP1-10-SmBiT or GFP11-LgBiT were mixed and seeded into 96-well plates. After 24 hours, cells were transfected with plasmids encoding various spike proteins using PEI. At 24 hours post-transfection, luciferase activity was measured using the Nano-Glo Luciferase Assay System (Promega #N1120), and luminescence was recorded with a GloMax luminometer (Promega). GFP fluorescence was detected using a FLUOstar Omega microplate reader (BMG LABTECH), and representative images were captured using an EVOS FL fluorescence microscope (Invitrogen).

### Western blot

To prepare cell lysates, cell pellets were resuspended in protein sample buffer (12.5% glycerol, 140 mM Tris-HCl, 2.5% SDS, 100 mM 2-mercaptoethanol, 0.5% bromophenol blue, pH 8.5), and lysed by sonication at 40% for 30 seconds (Branson #SFX150). After heating at 70 °C for 10 minutes, the samples were separated by electrophoresis on 4-12% Bis-Tris polyacrylamide gels (Thermo Fisher #NP0329BOX), followed by transfer of proteins to nitrocellulose membranes. Membranes were blocked with fish serum for 2 h at room temperature and probed with the primary antibodies at 4 °C overnight. After washing with TBST (100 mM NaCl, 10 mM Tris, pH 7.6, 0.1% Tween 20), blots were incubated with fluorescently-conjugated secondary antibodies for 1 h at room temperature, washed again with TBST, and scanned using a LiCor Odyssey scanner. To prepare pseudotyped particle samples for western blot analysis, the particles were concentrated by centrifugation through a sucrose cushion. Briefly, 1 mL of pseudotyped particle-containing supernatant was layered onto 250 μL of 20% sucrose and centrifuged at 20,000 × g for 16 h at 4 °C. The supernatant was carefully discarded, and the resulting pellet was resuspended in 50 μL of DMEM. Protein sample buffer was then added, and the samples were heated at 70 °C for 10 min before western blot analysis.

The antibodies used are as follows: rabbit anti-furin (Proteintech #18413-1-AP, 1:1,000), rabbit anti-TMPRSS2 (Abcam #ab109131, 1:2,000), mouse anti-CTSL (Thermo Fisher #BMS1032, 1:1,000), mouse anti-GAPDH (Proteintech #60004-1-Ig, 1:100,000), rabbit anti-HA (Proteintech #51064-2-AP, 1:5,000), rabbit anti-SARS-CoV-2-S1 (CST #56996, 1:1,000), rabbit anti-MERS-CoV spike (CST #41859, 1:1,000), rabbit anti-MHV S1 (Creative Diagnostics #DPAB-CS24652, 1:1,000) and home-made mouse anti-HIV p24 (1:100). The fluorescently-conjugated secondary antibodies include goat anti-mouse (Thermo Fisher # SA5-10176, 1:10,000) and goat anti-rabbit (Thermo Fisher #35568, 1:10,000).

### RNA extraction, reverse transcription, and qPCR

Pseudotyped particles RNA was extracted using Total RNA Kit I (Omega #R6834-01) according to the manufacturer’s instructions. RNA was reverse transcribed into cDNA with the HiScript IV RT SuperMix for qPCR kit (Vazyme #R423-01). qPCR was performed using a SYBR Green-based detection method with the Taq Pro Universal SYBR qPCR Master Mix (Vazyme # Q712-02) on a QuantStudio 7 Flex Real-Time PCR System (Thermo Fisher). Melt curve analysis was performed at the end of each run and showed a single specific melting peak, indicating specific amplification without detectable nonspecific products. The number of pseudotyped particles genome copy were quantified by absolute RT-qPCR using a standard curve generated from serial dilutions of CSGW reporter plasmid. Viral genome copy numbers were quantified using primers targeting GFP. The primer sequences were as follows: GFP forward, 5′-AAGGGCATCGACTTCAAGG-3′; GFP reverse, 5′-TGCTTGTCGGCCATGATATAG-3′.

### Inhibitor treatment

During pseudotyped particle packaging, the medium was replaced 12 hours post-transfection with fresh medium containing a furin inhibitor Decanoyl-RVKR-CMK (Tocris Bioscience #3501) at the concentrations in the figure. At 48 hours post-transfection, the viral supernatants were collected and used for titer determination.

A549-ACE2 cells were seeded into 96-well plates and pretreated 2 hours with E64D (MCE # HY-100229), an inhibitor that blocks the entry pathway of the endosomal membrane fusion. Subsequently, 100 µL of SARS-CoV-2 pseudotyped particles produced either by the conventional or the high-efficiency platform was overlaid directly onto the cells. After 48 hours of infection, cells were fixed and the percentage of GFP-positive cells was analyzed by flow cytometry.

### Neutralisation assays

24 hours before transduction, A549-ACE2 cells were seeded into 96-well plates. Anti-SARS-CoV-2 neutralizing antibody (Sino Biological # 40150-D001) was three-fold serial diluted in DMEM, then mixed with 100 µL SARS-CoV-2 pseudotyped particles produced in furin-knockout 293T cells. The pseudotyped particles mixture were incubated at 37 °C for 1 h and added into A549-ACE2 cells. After 48 hours of infection, cells were fixed and the percentage of GFP-positive cells was analyzed by flow cytometry.

### Statistical analysis

Statistical significance was assigned when *P* values were <0.05 using Prism Version 10.5 (GraphPad). Data analysis was determined using ANOVA or unpaired t-test, depending on data distribution and the number of comparison groups. Figure 2A and 6 were partially created with BioRender.com.

## ACKNOWLEDGMENTS

This work was funded by a UKRI Horizon Europe Guarantee (Frontier Research Guarantee) award to SJR (EP/Y011414/1 for an ERC Starting Grant) and Medical Research Council (https://www.ukri.org/councils/mrc) awards MR/P022642/1 (SJW, SJR), MR/V01157X/1 (SJW, SJR).

## SUPPLEMENTARY FIGURES LEGENDS

**Supplementary Fig 1. Assessment of knockout efficiency. A.** Following lentiviral transduction and antibiotic selection, cells lines were analyzed by immunoblotting for protease expression, with GAPDH as a loading control. Numbers below the blots indicate relative band intensity quantified by ImageJ, normalized to GAPDH, and expressed relative to the control. **B.** Genomic DNA was extracted from the mixed population of TMPRSS2-targeted cells. The sgRNA-targeted regions were amplified by PCR, and editing efficiency was assessed using ICE analysis. **C.** Furin-knockout single-cell clone #7 was expanded, and genomic DNA was extracted. The sgRNA-targeted region was amplified by PCR and analyzed by nanopore sequencing. The resulting sequencing reads were processed using CRISPResso2 to characterize the editing profile.

**Supplementary Fig 2. Dominant negative experiment. A-B.** Pseudotyped particles were generated by co-transfecting wild-type and S-GSAS spike-expression plasmids at the indicated proportions, while keeping the total amount of spike plasmid. Infectious titer of MHV-JHM pseudotyped particles were measured in 293T-MmCEACAM1 cells (A), and titer of SARS-CoV-2 pseudotyped particles were measured in 293T-HsACE2 cells (B).

## SUPPLEMENTARY TABLE LEGENDS

Supplementary Table 1. sgRNA sequences of genes selected for CRISPR-Cas9 editing experiments (see Excel file).

